# Impact of Kuntai Capsules on LIF, IGF-1 and EGF Expression in the Implantation Window of Endometrium in Mice

**DOI:** 10.1101/507509

**Authors:** Li Tan, Xiao-Xu Li, Xi-Ying Chu, Yan Li, Li-Jing Wan

## Abstract

**Objective:** This study aims to investigate the influence of Kuntai capsules on the expression level of leukemia inhibitory factor (LIF), insulin-like growth factor-I (IGF-1) and epidermal growth factor (EGF) during the mouse’s implantation window of superovulation period and controlled ovarian hyperstimulation period.

**Methods:** A total of 90 female mice were randomly divided into six groups (n=15, each group). The RNA expression of EGF, LIF and IGF-1 in the endometrium on the 4^th^ day of pregnancy was detected using the fluorescence quantitative PCR method, and the relative expression of these three factors in each group was compared.

**Results:** The mRNA expression of these three factors in the endometrium during the mouse’s implantation window period was significantly lower in the superovulation group and COH group than in the control group (P<0.001), and was obviously lower in the superovulation group than in the COH group (P<0.05). The mRNA expression of these three factors in the endometrium remained obviously lower in the superovulation plus kuntai capsule group and COH plus kuntai capsule group, compared with the control group (P<0.01). The mRNA expression of these three factors in the endometrium was lower in control group than in the NS plus kuntai capsule group (P<0.05). Furthermore, the expression in the superovulation and COH groups was lower than in the COH plus kuntai capsule group (P<0.05).

**Conclusions:** Both the superovulation and COH procedure could reduce the mouse’s endometrial receptivity. The application of the Chinese medicine Kuntai capsule could partially improve a mouse’s endometrial receptivity through the increasing expression of three relevant cytokines during the implantation window.

## Introduction

Along with social development and economic growth, the number of people with infertility continues to increase year by year due to a variety of factors, including the worsening air pollution, heightened life stress, and obesity [1–4]. Against this background, Assisted Reproductive Technology (ART) has gradually caught the attention of infertility patients. Although ART has been around for over 30 years, it has not yet gained popularity due to low clinical pregnancy rate [5–6]. According to relevant literatures, 60% of all failures of *in vitro* fertilization-embryo transfers (IVF-ETs) are attributed to diminished endometrial receptivity [7]. Therefore, the key for increasing embryo implantation rate and clinical pregnancy rate is to improve endometrial receptivity. Multiple studies have shown that leukemia inhibitory factor (LIF), insulin-like growth factor-I (IGF-1), and epidermal growth factor (EGF) exhibit high expression levels during the implantation window of several different species of mammals [8,9]. In addition, the concerted effects of these factors contribute to the establishment of endometrial tolerance. Therefore, these cytokines are generally considered as markers for testing endometrial tolerance. However, the drugs commonly used to improve endometrium tolerance are still subject to a variety of problems, and it remains questionable whether aspirin and aromatase inhibitors can actually improve the rate of embryo implantation. The price of growth hormone is too high, and Chinese medicine has the disadvantage of patient compliance. In order to overcome these problems, there is a need to determine a safe, convenient and inexpensive drug that could improve endometrial receptivity.

Kuntai capsules have been applied to improve female ovarian function in clinic for several years. It has been reported that endometrial thickness [10] and endometrial receptivity [11] can be improved. In the present study, quantitative fluorescence PCR (QF-PCR) was used to analyze the effects of Kuntai capsules on the expression levels of three cytokines related to endometrial tolerance during the superovulation cycle, natural cycle and controlled ovarian hyperstimulation mice implantation stage, providing a theoretical basis for the clinical application of ART, improving a patient’s endometrial tolerance, and improving the clinical pregnancy rate.

## 1. Materials and Methods

### 1.1 General materials

Healthy clean-grade 8-10 week old Kunming (KM) mice (20-30 grams in body weight) were chosen for the present study. Mice housing conditions are as follows: room temperature, 18-25°C; humidity, 60%-70%; 12 hour illumination followed by 12 hour darkness; male and female mice are kept in separate cages; mice were given unlimited access to food and water. Female mice that appeared normal in two successive estrous cycles were used in the experiments. All mice were purchased from the Laboratory Animal Center of Zhenzhou University (Permit #: SCXK[Henan] 2010-0002).

### 1.2 Methods

#### 1.2.1 Technique for observing the estrous cycles of mice

For judging the estrous cycle of female mice, vaginal smears were obtained at 9 AM every day with medical cotton swabs for morphology observation and the classification of the shedding of vaginal cells. The specific operation procedures were as follows: The tail was lifted with the left hand to fully expose the vaginal orifice. Then, the features of the external genital were observed, a medical cotton swab pre-dipped in physiological saline solution was held with the right hand, and inserted it into the mouse’ vagina at 0.5 cm in depth. Next, the swab was slowly twirled, pulled out, and the secretion stuck on the swab was smeared on a glass slide. Then, the slide was place on the bench for one minute, fix in 95% ethanol for 15-20 minutes, and hematoxylin and eosin (H&E) staining was performed (hematoxylin for one minute, eosin for 50 seconds). The slide was washed with distilled water, dried out, and observed under a microscope to ascertain the cell morphology. The mice estrous cycle can be divided into four phases, according to the features of the external genitals and vaginal smears of mice:

1. Proestrus: The vulva is pink, the vagina orifice opens up, and the periphery is swollen with serous fluid. As observed in the vaginal smear, there are less vaginal epithelial cells, the cytoplasmic stain is uneven, and few nucleated keratinocytes and an abundant number of leukocytes can be seen.
2. Estrus: The vulva is swollen and damp, the vaginal orifice is significantly enlarged and gray white, and the mucosa is almost dry. The vaginal smear shows that almost only nuclear keratinocytes can be seen.
3. Metestrus: The vulva is not significantly swollen, and the vagina orifice is closed or partially closed, and is light white in color. The vaginal smear shows that the keratinized epithelial cells are in clusters or swathes, and some leukocytes can be seen at the same time.
4. Anestrus: There is no significant change in the vulva, and the vaginal orifice is closed. The vaginal smear shows that the epithelial cells are small and shriveled, with large amounts of white blood cells in the smear.

#### 1.2.2 Experiment Group and Modeling

Female mice with two successive normal estrous cycles are divided into six groups, with 15 mice in each group: control group, saline solution plus Kuntai capsule group, superovulation group, superovulation plus Kuntai capsule group, controlled ovarian hyperstimulation (COH) group, and COH plus Kuntai capsule group. The animal model was established as described below.

1. Control group: At 9 AM every day, 0.3 ml of saline solution was applied on each mouse *via* intraperitoneal injection for nine days in succession. At the 9^th^ day, an extra saline solution was injected at 0.3 ml per mouse. After 48 hours, 0.3 ml was applied per mouse *via* intraperitoneal injection. Each female mouse was paired one-on-one with a male after ovulation. The appearance of the vaginal plug was check at 6 AM of the next day. Mice with a vaginal plug was marked as pregnancy day one and feed with 0.3 ml of saline solution per day until euthanization.
2. Saline solution plus Kuntai capsule group: At 9 AM every day, 0.3 ml of saline solution was applied per mouse *via* intraperitoneal injection for nine days in succession. At the 9^th^ day, extra saline solution was injected at 0.3 ml per mouse. After 48 hours, 0.3 ml was applied per mouse *via* intraperitoneal injection. Each female mouse was paired one-on-one with a male after ovulation. The appearance of the vaginal plug was check at 6 AM of the next day. Mice with a vaginal plug was marked as pregnancy day one and feed with 90.1 mg/100 g of body weight of solution of Kuntai capsule powder per day until euthanization.
3. Superovulation group: At 9 AM every day, 0.3 ml of saline solution was applied per mouse *via* intraperitoneal injection for nine days in succession. At the 9^th^ day, 10 IU of PMSG was injected per mouse. After 48 hours, 10 IU of HCG was applied per mouse *via* intraperitoneal injection. Each female mouse was paired one-on-one with a male during the evening. The appearance of the vaginal plug was checked at 6 AM of the next day. Mice with a vaginal plug were marked as pregnancy day one and fed with 0.3 ml of saline solution per day until euthanization.
4. Superovulation plus Kuntai capsule group: At 9 AM every day, 0.3 ml of saline solution was applied per mouse *via* intraperitoneal injection for nine days in succession. At the 9^th^ day, 10 IU of PMSG was injected per mouse. After 48 hours, 10 IU of HCG was applied per mouse *via* intraperitoneal injection. Each female mouse was paired one-on-one with a male during the evening. The appearance of the vaginal plug was checked at 6 AM of the next day. Mice with a vaginal plug was marked as pregnancy day one, and feed with 90.1 mg/100 g of body weight of solution of KUNTAI Capsule powder per day until euthanization.
5. COH group: At the beginning of metestrus, 1.5 ug/100 grams of body weight of GnRHa was applied per mouse at 9 AM everyday *via* intraperitoneal injection for nine days in succession. At the 9^th^ day, 10 IU of PMSF was injected per mouse. After 48 hours, 10 IU of HCG was applied per mouse *via* intraperitoneal injection. Each female mouse was paired one-on-one with a male during the evening. The appearance of the vaginal plug was checked at 6 AM of the next day. Mice with a vaginal plug was marked as pregnancy day one and feed with 0.3 ml of saline solution per day until euthanization.
6. COH plus Kuntai capsule group: At the beginning of metestrus, 1.5 ug/100 grams of body weight of GnRHa was applied per mouse at 9 AM everyday *via* intraperitoneal injection for nine days in succession. At the 9^th^ day, 10 IU of PMSF was injected per mouse. After 48 hours, 10 IU of HCG was applied per mouse *via* intraperitoneal injection. Each female mouse was paired one-on-one with a male during the evening. The appearance of the vaginal plug was checked at 6 AM of the next day. Mice with a vaginal plug were marked as pregnancy day one and fed with 90.1 mg/100 g of body weight of solution of Kuntai capsule powder per day until euthanization.

At 9 AM of the fourth day after pregnancy, anesthetics were applied to all mice *via* intraperitoneal injection. Caesarean section was performed to remove the uterus. A piece of endometrial tissue was slice off from the uterus on ice and weighed on an electronic scale. The tissue was immediately transferred to a centrifuge tube with 1 ml of Trizol lysis buffer. An electronic homogenizer was used to homogenize the tissues for 15 seconds in an ice-bath, and the tissue sample was stored in a freezer at −80°C.

#### 1.2.3 Detection Method of Quantitative Fluorescence PCR (QF-PCR)

Following the manual of the Trizol reagent, total RNA was extracted from mice endometrial tissues and reverse-transcribed into cDNA. The reactions were set up on ice. The system of the reverse transcription were RT Master Mix (2×) at 10 μl/l, RT Enzyme Mix at 1 μl/l, Oligo (dT) Random at 1 μl/l, specific primer (1 μM) at 1.2 μl/l, RNA template at 5-8 μl/l, RNase Free ddH_2_O, and total reaction at 20 μ. The conditions of the reverse-transcrption were 45°C for 40 minutes, and 85°C for 10 minutes.

LIF, IGF-1, EGF and β-actin (internal control) are amplified using cDNA reverse-transcribed from RNA as the template. The full DNA sequences of LIF, IGF-1, EGF and β-actin were obtained by searching the Genbank (Table 1).

**Table 1.**
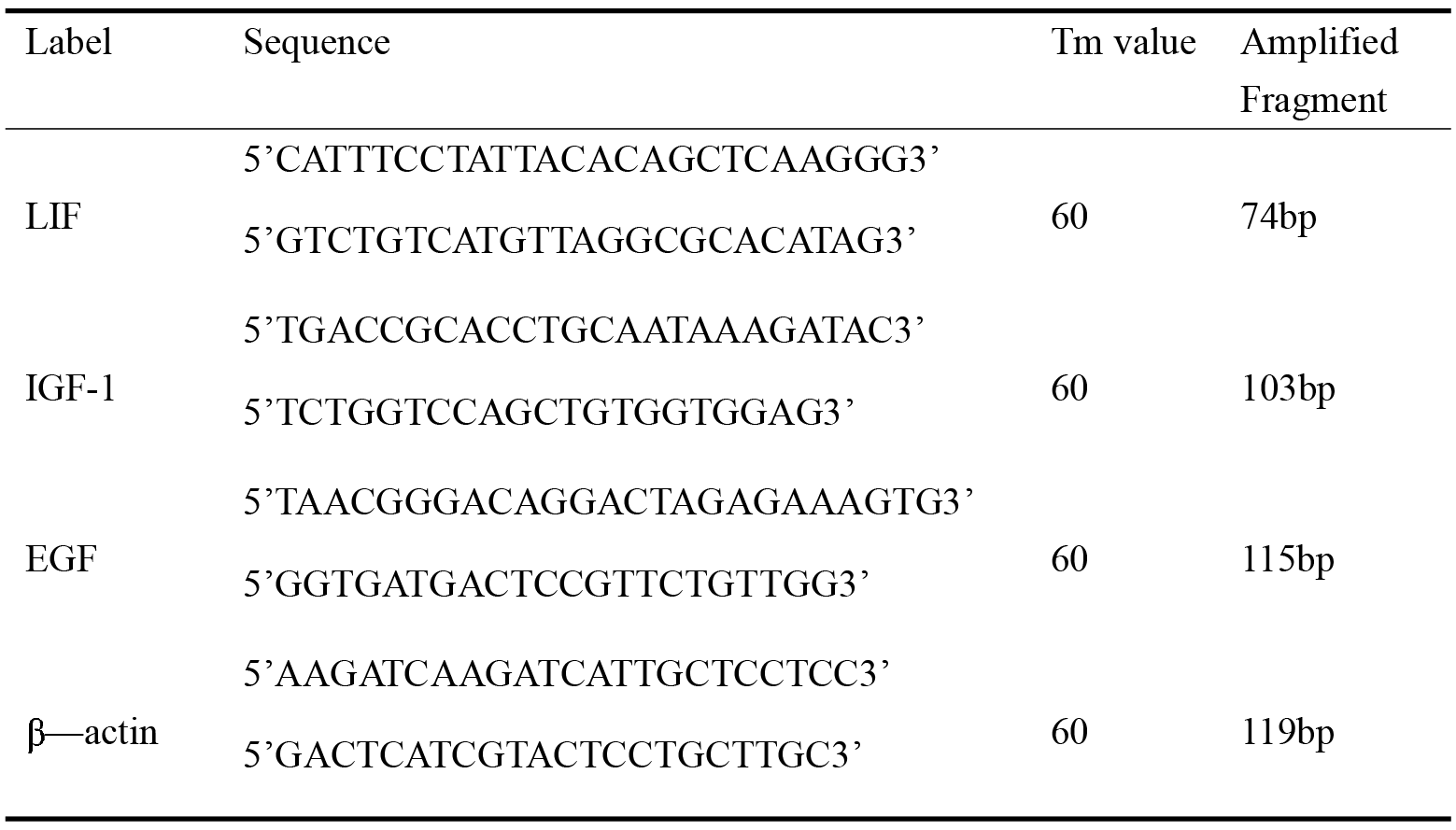
Primer Sequence

The reaction composition were cDNA template at X μl, forward primer (5 μM) at 0.4 μl, reverse primer (5 μM) at 0.4 μl, PCR Master Mix (2×) at 10 μl, ddH_2_O, and total reaction at 20 μ. The conditions were 94°C for two minutes, 94°C for 15 seconds, and 62°C for 40 seconds, for a total of 40 cycles.

When analyzing the fluorescence intensities of each sample with QF-PCR, the relative quantitative ^□□^CT method was used to detect changes in gene expression. Before the ^□□^CT test, a pilot experiment was performed to validate its feasibility. An easy dilution was used to prepare serial dilutions of one sample cDNA (1×, 10×, 100×, 1,000×, 10,000×). QF-PCR assay was conducted on these dilutions with β-actin as the internal control. The ^□□^CT was only applicable if it was under the same reaction composition and conditions, the difference between the amplification curve slopes of the target gene and internal control was <0.1 [12].

### 1.3 Statistical Analysis

All data were processed using SPSS 17.0. The quantitative comparison of inter-group data was achieved using *t*-test. Univariate analysis was used for quantitative comparisons between multiple sets of data. LSD *t*-test was applied to achieve one-on-one comparisons among multiple sets of data. A P-value <0.05 was considered statistically significant.

## 2. Results

### 2.1 Electrophoresis results of LIF, IGF-1 and EGF in all groups

#### 2.1.1 Electrophoresis results of LIF in all groups (Fig. 1)

**Figure 1.**
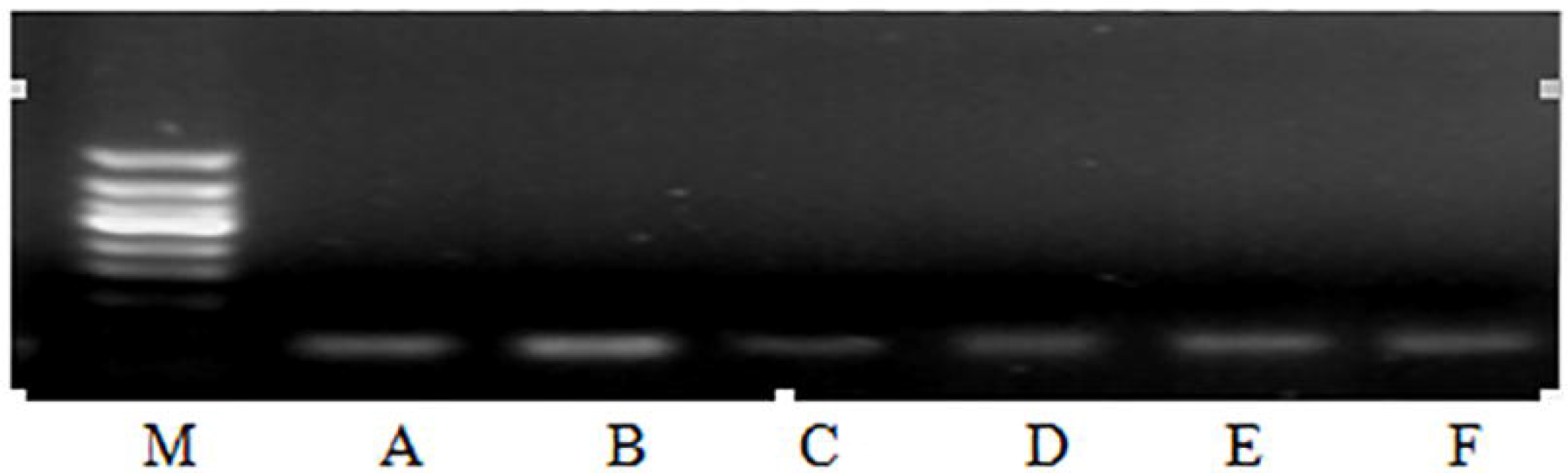
The electrophoresis results of LIF in group A~F

#### 2.1.2 Electrophoresis results of IGF-1 in all groups (Fig. 2)

**Figure 2.**
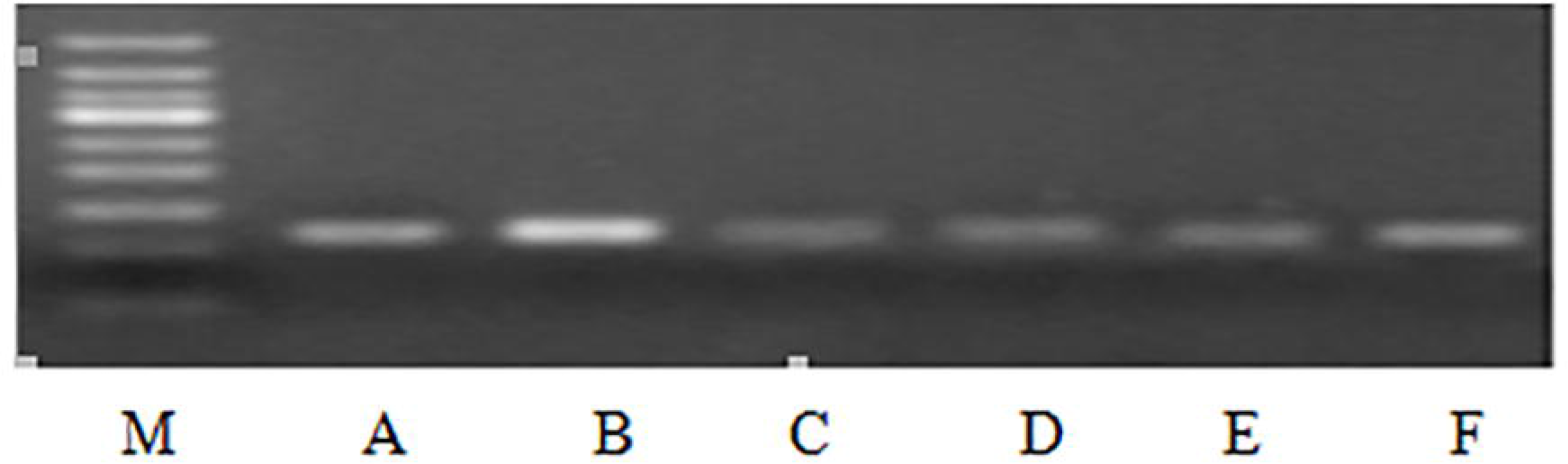
The electrophoresis results of IGF-1 in group A~F

#### 2.1.3 Electrophoresis results of EGF in all groups (Fig. 3)

**Figure 3.**
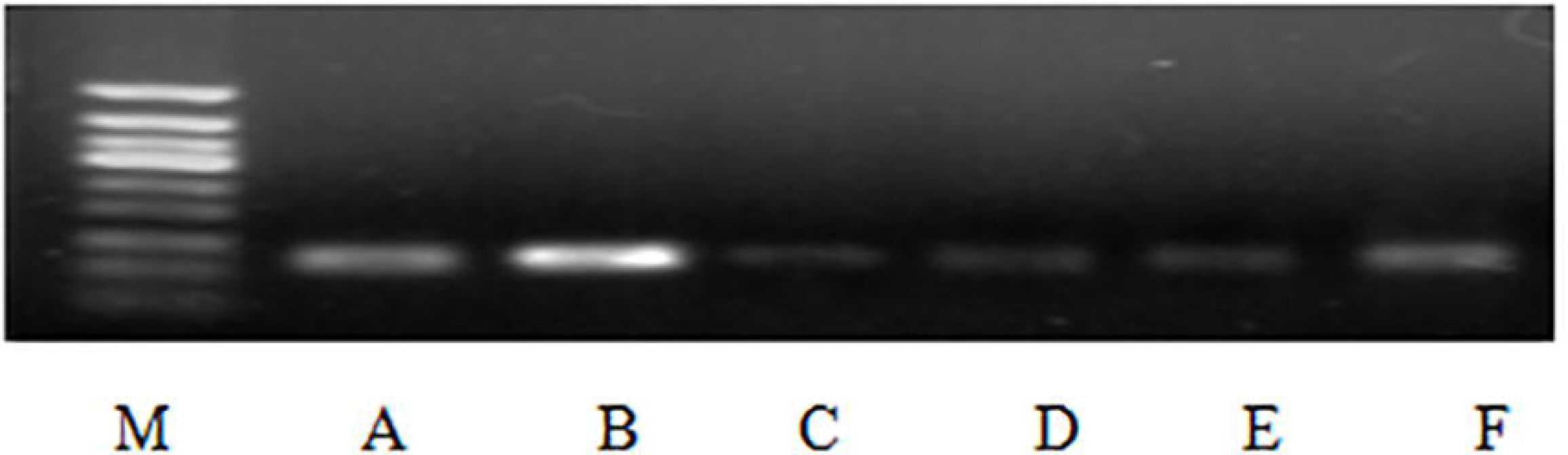
The electrophoresis results of EGF in group A~F

### 2.2 The expression levels of LIF, IGF-1 and EGF in the different groups

#### 2.2.1 The expression of LIF, IGF-1 and EGF in mouse entrometrial tissues in the control group and saline solution plus Kuntai capsules group

The mRNA levels of LIF, IGF-1 and EGF were high in endometrial tissues in the control group and saline solution plus Kuntai capsule group at the implantation window phase. QF-PCR results also suggest that all three genes exhibited higher expression levels in the saline solution plus Kuntai capsules group than in the control group. The differences were statistically significant, as suggested by the mean *P*-value (<0.001) (Table 2).

**Table 2.**
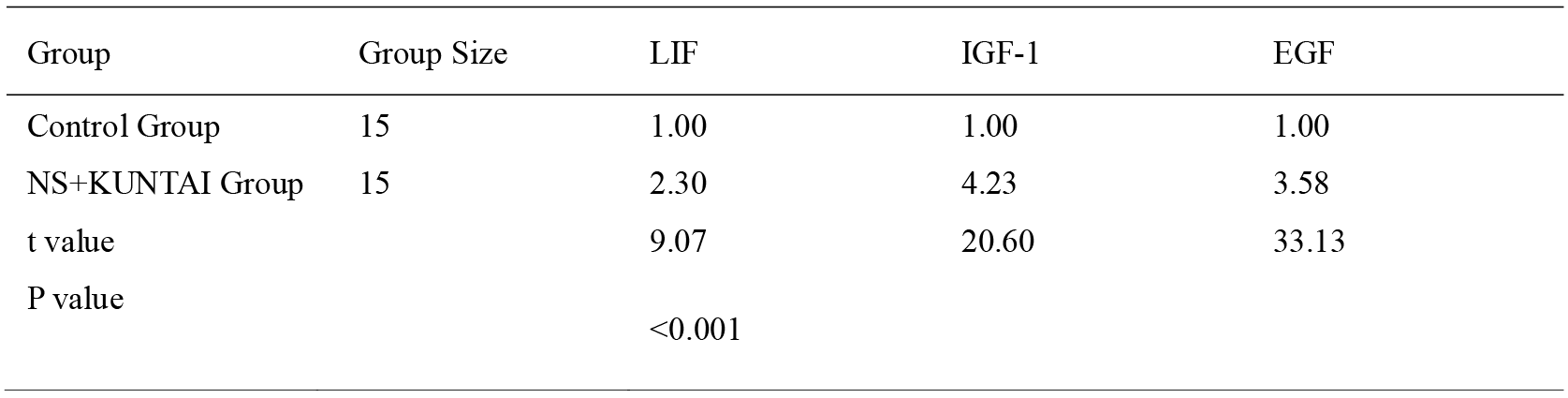
Comparison of relative expression levels of LIF, IGF-1, EGF mRNA in control group
and NS+KUNTAI Capsule group

#### 2.2.2 The expression of LIF, IGF-1 and EGF in murine entrometrial tissues in the superovulation group and superovulation plus Kuntai capsule group

The mRNA levels of LIF, IGF-1 and EGF were low in endometrial tissues in the superovulation group and superovulation plus Kuntai capsule group at the implantation window phase. However, QF-PCR results also suggest that all three genes exhibited higher expression levels in the superovulation plus Kuntai capsule group than in the superovulation group. The differences were statistically significant, as suggested by the mean *P*-value (<0.001) (Table 3).

**Table 3.** Comparison of relative expression levels of LIF, IGF-1, EGF mRNA in Superovulation
group and Superovulation+KUNTAI Capsule group

| Group | Group Size | LIF | IGF-1 | EGF |
| --- | --- | --- | --- | --- |
| Superovulation Group | 15 | 1.00 | 1.00 | 1.00 |
| Superovulation+KUNTAI Group | 15 | 8.45 | 3.32 | 5.90 |
| t value |  | 26.20 | 28.38 | 27.27 |
| P value |  | P<0.001 |  |  |

#### 2.2.3 The expression of LIF, IGF-1 and EGF in murine entrometrial tissues in the COH group and COH plus Kuntai capsule group

The mRNA levels of LIF, IGF-1 and EGF were low in endometrial tissues in the COH group, while medium levels were observed in samples obtained from the COH plus Kuntai capsule group. QF-PCR results suggest that all three genes exhibited higher expression levels in the COH plus Kuntai capsule group than in the COH group. The differences were statistically significant, as suggested by the mean *P*-value (<0.001) (Table 4).

**Table 4.**
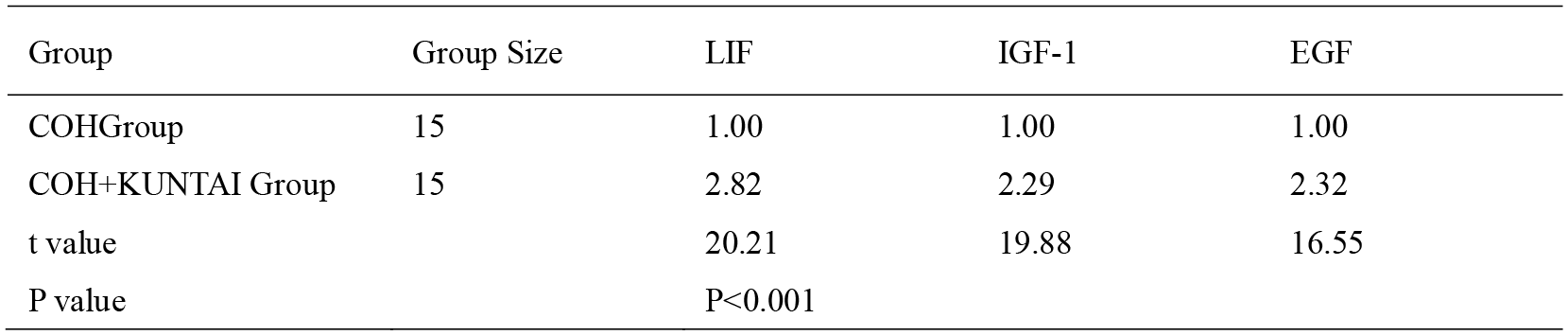
Comparison of relative expression levels of LIF, IGF-1, EGF mRNA in COH group and
COH+KUNTAI Capsule group

### 2.3 Effects of superovulation and COH on the expression levels of LIF, IGF-1 and EGF in murine endometrial tissues

The mRNA levels of LIF, IGF-1 and EGF were significantly higher in the control group than in the superovulation group (mean *P*-value <0.001) and COH group, and the differences were statistically significant (mean *P*-value <0.001). Moreover, these levels in the COH group were higher than in the superovulation group, and the differences were also statistically significant (mean *P*-value <0.001) (Table 5).

**Table 5.**
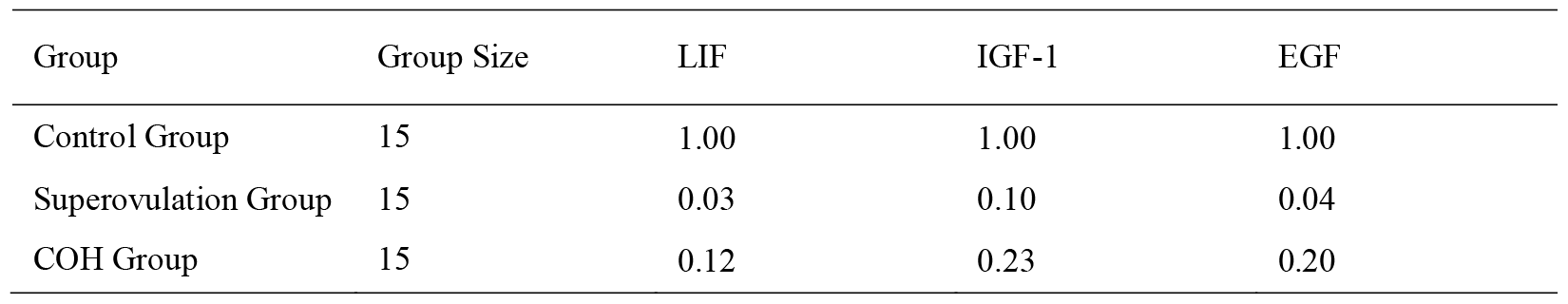
The relative mRNA expression levels of three cytokines in control group,
Superovulation group and COH group with group A as the reference

### 2.4 Effects of Kuntai capsules on the expression levels of LIF, IGF-1 and EGF in murine endometrial tissues

The mRNA levels of LIF, IGF-1 and EGF in the control group were markedly higher than in the superovulation plus Kuntai capsule group and COH plus Kuntai capsule group, and the differences were statistically significant, as suggested by mean *P*-values <0.001. In addition, all three genes exhibited a higher mRNA expression in the COH plus Kuntai capsule group than in the superovulation plus Kuntai capsule group, and the difference was statistically significant (mean *P*-value <0.01) (Table 6).

**Table 6.** The relative mRNA expression levels of three cytokines in control group,
Superovulation+KUNTAI group and COH+KUNTAI group with group A as the reference

| Group | Group Size | LIF | IGF-1 | EGF |
| --- | --- | --- | --- | --- |
| Control Group | 15 | 1.00 | 1.00 | 1.00 |
| Superovulation+KUNTAI Group | 15 | 0.27 | 0.34 | 0.26 |
| COH+KUNTAI Group | 15 | 0.35 | 0.52 | 0.47 |

## 3. Discussion

Endometrial tolerance refers to the endometrium’s capability to receive embryos, which is crucial for a successful embryonic implantation. The endometrial process of embryonic implantation and development is completed within a short period of time, which is called the implantation window or nidation window. Numerous studies have confirmed that the establishment of endometrial tolerance during the implantation window is mediated through the concerted effects of multiple cytokines. As the most representative ones among these cytokines, LIF, IGF-1 and EGF play crucial roles in regulating endometrial tolerance. As a means to promote the simultaneous development of multiple follicles, superovulation and COH have become an important part of ART therapy. Nonetheless, according to related studies [13,14], superovulation and COH may disrupt the synchrony between embryonic development and the opening of the implantation window, which would lead to diminished endometrial tolerance and *in vitro* fertilization (IVF) pregnancy rate. Therefore, we need to identify a therapeutic strategy that could minimize the impact of superovulation on endometrial tolerance.

Medicines presently used for this purpose include low-dose aspirin, aromatase inhibitor, growth hormone and Chinese Medicines such as donkey-hide gelatin. However, the actual efficacy of aspirin and aromatase inhibitor in improving embryonic implantation and clinical pregnancy remains controversial. The usage of growth hormone is limited due to its high cost, whereas Chinese medicine has a number of drawbacks such as inconvenient intake and poor patient adherence. In order to overcome these challenges, there is a need to seek for a safe, convenient and inexpensive medicine that could improve endometrial tolerance. Kuntai capsule is a formulated traditional Chinese medicine that comprise of six herb components: *Rehmannia glutinosa* as the primary ingredient, *Radix Paeoniae Alba, Coptis chinensis* and *donkey-hide gelatin* as the secondary ingredients, and *Scutellaria baicalensis* and *Poria cocos* as the adjuvant ingredients. By maintaining the Yin and Yang balance, it is supposed to achieve the efficacies of supplementing the kidney, nurturing the liver, nourishing the Yin and dousing body flames, and soothing the nerves. Multiple studies [15–18] have suggested that Chinese medicine for nourishing the kidney and invigorating blood circulation could improve the blood circulation of the endometrim to a certain extent. By promoting blood supply to endometrium and increasing endometrium thickness and the number of glands, such medicines can help increase the clinical pregnancy rate.

In the present study, QF-PCR technology was employed to quantitatively evaluate the effects of Kuntai capsules on endometrial tolerance. Results revealed that the mRNA expression levels of the three cytokines are markedly higher in the saline solution plus Kuntai capsules group than in the control group. Moreover, these expression levels were higher in the superovulation plus Kuntai capsule group and COH plus Kuntai capsule group, compared to the superovulation group and COH group. All differences were statistically significant (mean *P*-value <0.001), indicating the efficacy of Kuntai capsules in improving murine endometrial tolerance. Nonetheless, based on the univariate analysis of the mRNA levels of these three cytokines in the control group, superovulation plus Kuntai capsule group and COH plus Kuntai capsule group, the control group exhibited the highest levels, followed by the superovulation plus Kuntai capsule group and COH plus Kuntai capsule group. This indicates that despite the positive effect of Kuntai capsules on murine endometrial tolerance, it is insufficient to offset damages to the endometrium induced by superovulation and COH. Chu Xiying *et al.* [19] evaluated the effects of Kuntai capsules on murine endometrial thickness through immunohistochemistry, and demonstrated that Kuntai capsules can significantly enhance the endometrial protein levels of LIF and IGF-1. Moreover, they revealed that these levels were significantly higher in the superovulation plus Kuntai capsule group and COH plus Kuntai capsule group, compared to the superovulation group and COH group. Another study [20] demonstrated that high doses of Kuntai capsules did not display early embryonic toxicity to SD rats, and does not affect the reproductivity of rats, suggesting of the superb safety of Kuntai capsules in clinical application. Thus, in our future work, we may consider increasing the dosage of Kuntai capsule or combining it with other medicines, with the hope of finding a more ideal drug that could neutralize endometrial damages caused by superovulation and COH, and increase the clinical pregnancy rate of IVF-ET.

In summary, the application of Chinese medicine Kuntai capsule in IVF-ET can partially improve murine endometrial tolerance by increasing the expression of three relevant cytokines during the implantation window. However, it is insufficient to completely reverse the endometrial damages caused by superovulation and COH. In the present study, only conventional doses of Kuntai capsule power solutions were tested. Since previous studies have demonstrated the safety of large doses of Kuntai capsules, we can identify the maximum remedial effect of Kuntai capsules on damaged endometrial tolerance in the future, providing reliable experimental evidences for further clinical research.

## References

[1] Gies, Erica. Even Conscientious People Have An Eco-Footprint[J]. EN,2012,27(1): 51–52.

[2] Aliyeh G, Laya F. Quality of life and its correlates among a group of infertile Iranian women. Med Sci Monit 2007;13:CR313–317.

[3] Broughton DE, Moley KH. Obesity and Female Infertility: Potential Mediators of Obesity’s Impact. Fertil Steril 2017;107:840–847.

[4] Ding J, Luo XT, Yao YR, Xiao HM, Guo MQ. Investigation of changes in endocannabinoids and N-acylethanolamides in biofluids, and their correlations with female infertility. J Chromatogr A 2017;1509:16–25.

[5] Rhodes TL, McCoy TP, Higdon HL 3rd, Boone WR. Factors affecting assisted reproductive technology (ART) pregnancy rates: a multivariate analysis. J Assist Reprod Genet 2005;22:335–346.

[6] Yang R, Yang S, Li R, Chen X, Wang H, Ma C, Liu P, Qiao J. Biochemical pregnancy and spontaneous abortion in first IVF cycles are negative predictors for subsequent cycles: an over 10,000 cases cohort study. Arch Gynecol Obstet 2015;292:453–458.

[7] Moreno I, Franasiak JM. Franasiak.Endometrial microbiota—new player in town. Fertil Steril 2017;108:32–39.

[8] Pathare ADS, Zaveri K, Hinduja I. Downregulation of genes related to immune and inflammatory response in IVF implantation failure cases under controlled ovarian stimulation. Am J Reprod Immunol 2017;78(1). doi: 10.1111/aji.12679.

[9] Islam MR, Yamagami K, Yoshii Y, Yamauchi N. Growth factor induced proliferation, migration, and lumen formation of rat endometrial epithelial cells in vitro. J Reprod Dev 2016;62:271–278.

[10] Chu X, Song Y, Wan L, Tan L. Effects of kuntai capsule on endometrial thickness and the expression of epidermal growth factor proteins in the uterus of control superovulation mice. Zhonghua Yi Xue Za Zhi 2014;94:2300–2303.

[11] Wu CJ. Effects of kuntai capsule on follicular development, endometrium and ovulation in ovulatory infertility. China practical medicine 2015;(18):10–11.

[12] Livak KJ, Schmittgen TD. Analysis of relative gene expression data using real-time quantitative PCR and the 2(-Delta DeltaC(T)) Method. Methods 2001;25:402–408.

[13] Meyer WR, Novotny DB, Fritz MA, Beyler SA, Wolf LJ, Lessey BA. Effect of exogenous on endometrial maturation in oocyte donors. Fertil Steril 1999;71:109–114.

[14] Chillik C, Acosta A. THE role of LHRH agonists and antagonists. Reprod Biomed Online 2001;2:120–128.

[15] Zhang SC, Wu ZK, Cai LX, Shen MX. Theoretic significance of “promoting sexuality through nurturing kidney” based on the experimental evidences for improving effects of kidney nourishing Chinese medicine on tissue angiogenesis. Study Journal of Traditional Chinese Medicine 2005;23:1078–1080.

[16] Wang WJ. Clinical and experimental studies on treating embryonic implantation failures with Chinese medicines for nourishing kidney and limbs [Master Thesis]. Wuhan: Hubei University of Chinese Medicine, 2010.

[17] Huang DM, Huang GY, Lu FE. Impacts on blastocyst implantation dysfunction and murine endometrial glands apoptosis by Chinese medicines for nourishing kidney, soothing nerves and improving blood circulation. Journal of Traditional Chinese Medicine 2008;49:644–646.

[18] Nan Y, Duan YX, Li YJ. Impacts of KUNTAI Capsule on endometrial tolerance of infertility patients. Journal of Xinxiang Medical School 2012;29:384–385.

[19] Chu XY. The effects of Chinese Medicine KUNTAI Capsule on endometrial thickness, EGF, IGF-1, PCNA and LIF protein levels during murine reproductive phase. [Master Thesis]. Henan: Zhenzhou University No.2 Affiliated Hospital, 2013.

[20] Wang W, Zhang Y, Liu K, Gong HY, He JL, Gong L. Study on impacts on SD rats’ reproductivity and early embryonic toxicity of KUNTAI Capsule. Chinese Traditional Patent Medicine 2012;34:1869–1873.

